# Light Attention Predicts Protein Location from the Language of Life

**DOI:** 10.1101/2021.04.25.441334

**Authors:** Hannes Stärk, Christian Dallago, Michael Heinzinger, Burkhard Rost

## Abstract

**Summary:** Although knowing where a protein functions in a cell is important to characterize biological processes, this information remains unavailable for most known proteins. Machine learning narrows the gap through predictions from expert-designed input features leveraging information from multiple sequence alignments (MSAs) that is resource expensive to generate. Here, we showcased using embeddings from protein language models (pLMs) for competitive localization prediction without MSAs. Our lightweight deep neural network architecture used a softmax weighted aggregation mechanism with linear complexity in sequence length referred to as light attention (LA). The method significantly outperformed the state-of-the-art (SOTA) for ten localization classes by about eight percentage points (Q10). So far, this might be the highest improvement of *just embeddings* over MSAs. Our new test set highlighted the limits of standard static data sets: while inviting new models, they might not suffice to claim improvements over the SOTA.

**Availability:** Online predictions are available at http://embed.protein.properties. Predictions for the human proteome are available at https://zenodo.org/record/5047020. Code is provided at https://github.com/HannesStark/protein-localization.

## 1. Introduction

### Prediction bridges gap between proteins with and without location annotations

Proteins are the machinery of life involved in all essential biological processes (*Appendix: Biological Background*). Knowing where in the cell a protein functions, *natively*, i.e. its *subcellular location* or *cellular compartment* (for brevity, abbreviated by *location*), is important to unravel biological function (Nair & Rost, 2005; Yu et al., 2006). Experimental determination of protein function is complex, costly, and selection biased Ching et al., 2018. In contrast, protein sequences continue to explode (Consortium, 2021). This increases the sequence-annotation gap between proteins for which only the sequence is known and those with experimental function annotations. Computational methods have been bridging this gap (Rost et al., 2003), e.g. by predicting protein location (Goldberg et al., 2012; 2014; Almagro Armenteros et al., 2017; Savojardo et al., 2018). The standard tool in molecular biology, namely homology-based inference (<u>HBI</u>), accurately transfers annotations from experimentally annotated to sequence-similar un-annotated proteins. However, HBI is either unavailable or unreliable for most proteins (Goldberg et al., 2014; Mahlich et al., 2018). Machine learning methods perform less well (lower precision) but are available for all proteins (high recall). The best methods use evolutionary information as computed from families of related proteins identified in multiple sequence alignments (<u>MSAs</u>) as input (Nair & Rost, 2005; Goldberg et al., 2012; Almagro Armenteros et al., 2017). Although the marriage of evolutionary information and machine learning has influenced computational biology for decades (Rost & Sander, 1993), due to database growth, MSAs have become costly.

### Protein Language Models (<u>pLMs</u>) better represent sequences

Recently, protein sequence representations (embeddings) have been learned from databases (Steinegger & Söding, 2018; Consortium, 2021) using language models (<u>LMs</u>) (Bepler & Berger, 2019; Alley et al., 2019; Heinzinger et al., 2019; Rives et al., 2021; Elnaggar et al., 2021) initially used in natural language processing (<u>NLP</u>) (Peters et al., 2018; Devlin et al., 2019; Raffel et al., 2020). Models trained on protein embeddings via transfer learning tend to be outperformed by approaches using MSAs (Rao et al., 2019; Heinzinger et al., 2019). However, embedding-based solutions can outshine HBI (Littmann et al., 2021) and advanced protein structure prediction methods (Bhattacharya et al., 2020; Rao et al., 2020; Weißenow et al., 2021). Yet, for location prediction, embedding-based models (Heinzinger et al., 2019; Elnaggar et al., 2021; Littmann et al., 2021) remained inferior to the state-of-the-art (<u>SOTA</u>) using MSAs, such as DeepLoc (Almagro Armenteros et al., 2017).

In this work, we leveraged protein embeddings to predict cellular location without MSAs. We proposed a deep neural network architecture using light attention (LA) inspired by previous attention mechanisms (Bahdanau et al., 2015).

## 2. Related Work

The best previous predictions of location prediction combined HBI, MSAs, and machine learning, often building prior expert-knowledge into the models. For instance, *Loc-Tree2* (Goldberg et al., 2012) implemented profile-kernel (Support Vector Machines (<u>SVMs</u>) (Cortes & Vapnik, 1995) which identified k-mers conserved in evolution and put them into a hierarchy of models inspired by cellular sorting pathways. *BUSCA* (Savojardo et al., 2018) combined three compartment-specific SVMs based on MSAs (Pierleoni et al., 2006; Savojardo et al., 2017). *DeepLoc* (Almagro Armenteros et al., 2017) used convolutions followed by a bidirectional long short-term memory (<u>LSTM</u>) module (Hochreiter & Schmidhuber, 1997) employing the Bahdanau-Attention (Bahdanau et al., 2015). Using the BLOSUM62 substitution metric (Henikoff & Henikoff, 1992) for fast and MSAs for slower, refined predictions, DeepLoc rose to become the SOTA. Embedding-based methods (Heinzinger et al., 2019) have not yet consistently outperformed this SOTA, although *ProtTrans* (Elnaggar et al., 2021), based on very large data sets, came close.

## 3. Methods

### 3.1. Data

#### Standard setDeepLoc

Following previous work (Heinzinger et al., 2019; Elnaggar et al., 2021), we began with a data set introduced by *DeepLoc* (Almagro Armenteros et al., 2017) for training (13 858 proteins) and testing (2 768 proteins). All proteins have experimental evidence for one of ten location classes (nucleus, cytoplasm, extracellular space, mitochondrion, cell membrane, Endoplasmatic Reticulum, plastid, Golgi apparatus, lysosome/vacuole, peroxisome). The 2 768 proteins making up the test set (dubbed *setDeepLoc*), had been redundancy reduced to the training set (but not to themselves), and thus share ≤ 30% <u>PIDE</u> (pairwise sequence identity) and E-values ≤ 10^−6^ to any sequence in training. To avoid overestimations by tuning hyper-parameters, we split the DeepLoc training set into: training-only (9 503 proteins) and validation sets (1 158 proteins; ≤ 30% PIDE; *Appendix: Datasets*).

#### Novel *setHARD*

To catch over-fitting on a static standard data set, we created a new independent test set from *SwissProt* (Consortium, 2021). Applying the same filters as *DeepLoc* (only eukaryotes; all proteins ≥ 40 residues; no fragments; only experimental annotations) gave 5 947 proteins. Using *MMseqs2* (Steinegger & Söding, 2017), we removed all proteins from the new set with ≥ 20% PIDE to any protein in any other set. Next, we mapped location classes from DeepLoc to SwissProt, merged duplicates, and removed multi-localized proteins (protein X both in class Y and Z). Finally, we clustered at ≥ 20% PIDE leaving only one representative of each cluster in the new, more challenging test set (dubbed *setHARD;* 490 proteins; *Appendix: Datasets*).

### 3.2. Models

#### Input embeddings

As input to the Light Attention (<u>LA</u>) architectures, we extracted *frozen* embeddings from pLMs, i.e. without fine-tuning for location prediction (details below). We compared embeddings from five main and a sixth additional pre-trained pLMs (Table 1): (1) *<u>SeqVec</u>* (Heinzinger et al., 2019) is a bidirectional LSTM based on on ELMo (Peters et al., 2018) that was trained on UniRef50 (Suzek et al., 2015). (2) *<u>ProtBert</u>* (Elnaggar et al., 2021) is an encoder-only model based on BERT (Devlin et al., 2019) that was trained on BFD (Steinegger & Söding, 2018). (3) ProtT5-XL-UniRef50 (Elnaggar et al., 2021) (for simplicity: *<u>ProtT5</u>*) is an encoder-only model based on T5 (Raffel et al., 2020) that was trained on BFD and fine-tuned on Uniref50. (4) *<u>ESM-1b</u>* (Rives et al., 2021) is a transformer model that was trained on UniRef50. (5) *<u>UniRep</u>* (Alley et al., 2019) is a multiplicative LSTM (mLSTM)-based model trained on UniRef50. (6) Bepler&Berger (dubbed <u>*BB*</u>) is a bidirectional LSTM by (Bepler & Berger, 2019), which fused modelling the protein language with learning information about protein structure into a single pLM. Due to different training objectives, this pLM was expected suboptimal for our task. As results confirmed this expectation, we confined these to *Appendix: Additional Results*.

**Table 1.**
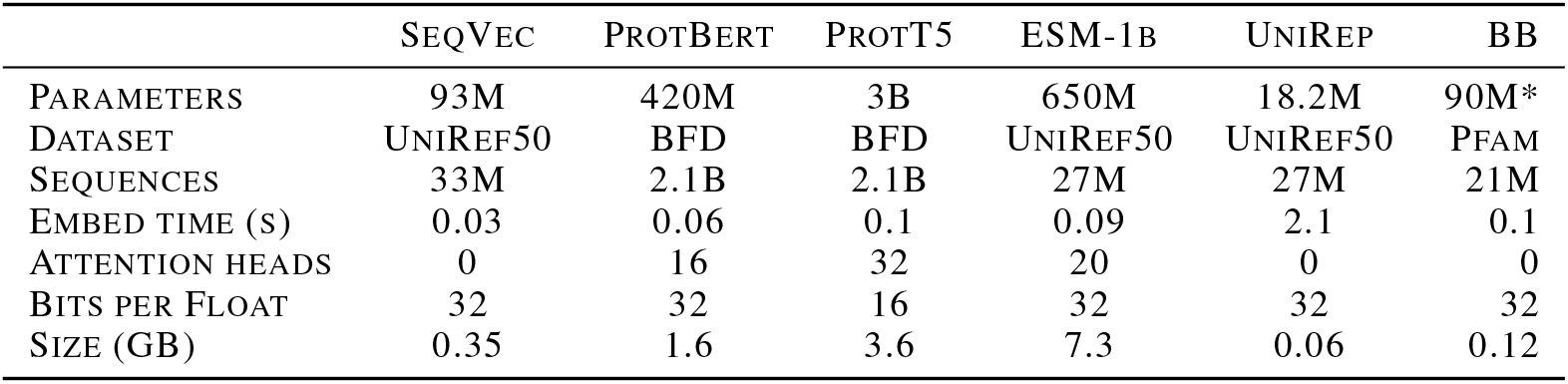
Implementation details for SeqVec (Heinzinger et al., 2019), ProtBert (Elnaggar et al., 2021), ProtT5 (Elnaggar et al., 2021), ESM-1b (Rives et al., 2021), UniRep (Alley et al., 2019) and BB (Bepler & Berger, 2019). Estimates marked by *; differences in the number of proteins (*Sequences*) for the same set (*Dataset*) originated from versioning. The embedding time (in seconds) was averaged over 10 000 proteins taken from the PDB (Berman et al., 2000) using the embedding models taken from *bio-embeddings* (Dallago et al., 2021).

Frozen embeddings were preferred over fine-tuned embeddings as the latter previously did not improve (Elnaggar et al., 2021) and consumed more resources/energy. *ProtT5* was instantiated at half-precision (float16 weights instead of float32) to ensure the encoder could fit on consumer GPUs with limited vRAM. Due to model limitations, for ESM-1b, only proteins with fewer than 1024 residues were used for training and evaluation (*Appendix: Datasets*).

Embeddings for each residue (NLP equivalent: word) in a protein sequence (NLP equivalent: document) were obtained using the bio-embeddings software (Dallago et al., 2021). For *SeqVec*, the per-residue embeddings were generated by summing the representations of each layer. For all other models, the per-residue embeddings were extracted from the last hidden layer. Finally, the inputs obtained from the pLMs were of size *d_in_* × *L*, where *L* is the length of the protein sequence, while *d_in_* is the size of the embedding.

#### Implementation details

The LA models were trained using filter size *s* = 9, *d_out_* = 1024, the Adam (Kingma & Ba, 2015) optimizer (learning rate 5 × 10^−5^) with a batch size of 150, and early stopping after no improvement in validation loss for 80 epochs. We selected the hyperparameters via random search (*Appendix: Hyperparameters*). Models were trained either on an Nvidia Quadro RTX 8000 with 48GB vRAM or an Nvidia GeForce GTX 1060 with 6GB vRAM.

#### Light Attention (LA) architecture

The input to the light attention (<u>LA</u>) classifier (Fig. 1) was a protein embedding 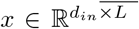 where *L* is the sequence length, while *d_in_* is the size of the embedding (which depends on the model, Table 1). The input was transformed by two separate 1D convolutions with filter sizes s and learned weights 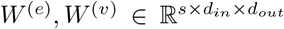. The convolutions were applied over the length dimension to produce attention coefficients and values *e*, 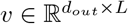

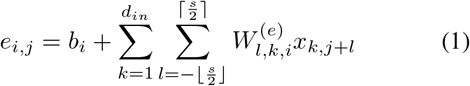

where 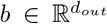 is a learned bias. For *j* ∉ [0, *L*), the *x*_:,*j*_ were zero vectors. To use the coefficients as attention distributions over all *j*, we *softmax-normalized them over the length dimension*, i.e. the attention weight 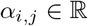 for the j-th residue and the i-th feature dimension was calculated as:

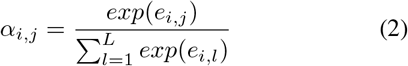

**Figure 1.**
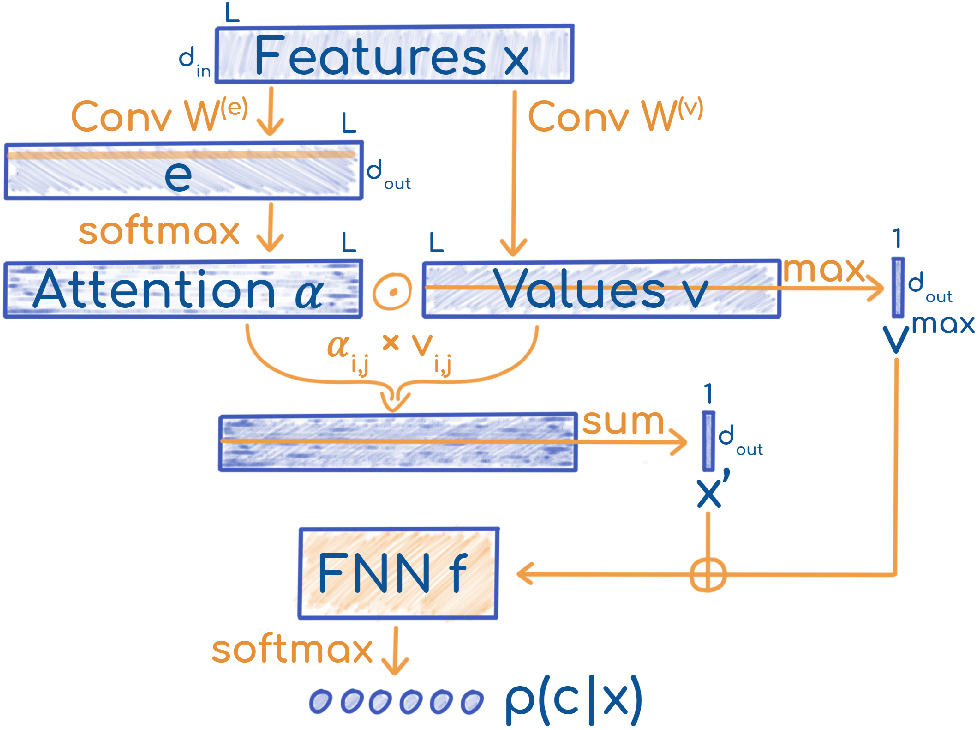
Sketch of Light Attention (LA). The LA architecture was parameterized by two weight matrices 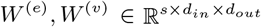 and the weights of an FNN 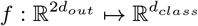.

Note that the weight distributions for each feature dimension *i* are independent, and they can generate different attention patterns. The attention distributions were used to compute weighted sums of the transformed residue embeddings *v_i,j_*. Thus, we obtained a fixed-size representation 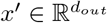 for the whole protein, independent of its length.

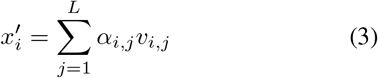

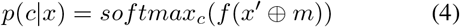

#### Methods used for comparison

For comparison, we trained a two-layer feed-forward neural network (<u>FNN</u>) proposed previously (Heinzinger et al., 2019). Instead of per-residue embeddings in 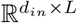, the FNNs used sequence-embeddings in 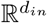, which derived from residue-embeddings averaged over the length dimension (i.e. mean pooling). Furthermore, for these representations, we performed embeddings distance-based annotation transfer (dubbed <u>EAT</u>) (Littmann et al., 2021). In this approach, proteins in *setDeepLoc* and *setHARD* were annotated by transferring the location from the nearest neighbor (L1 embedding distance) in the training set.

For ablations on the architecture, we tested LA without the softmax aggregation (<u>LA w/o Softmax</u>) that previously produced *x*′, by replacing it with averaging of the coefficients *e*. Then, with <u>LA w/o MaxPool</u>, we discarded the max-pooled values *v^max^* as input to the FNN instead of concatenating them with *x*′. With <u>Attention from *v*</u>, we computed the attention coefficients *e* via a convolution over the values *v* instead of over the inputs *x*. Additionally, we tested using a simple stack of convolutions (kernel-size 3, 9, and 15) followed by adaptive pooling to a length of 5 and an FNN instead of LA (<u>Conv + AdaPool</u>). Similarly, <u>Query-Attention</u> replaces the whole LA architecture with a transformer layer that used a single learned vector as query to summarize the whole sequence. As the last alternative operating on LM representations, we considered the <u>DeepLoc LSTM</u> (Almagro Armenteros et al., 2017) with *ProtT5* embeddings instead of MSAs.

To evaluate how traditional representations stack up against pLM embeddings, we evaluated MSAs (<u>LA(MSA)</u>) and one-hot encodings of amino acids (<u>LA(OneHot)</u>) as inputs to the LA model.

### 3.3. Evaluation

Following previous work, we assessed performance through the mean ten-class accuracy (Q10), giving the percentage of correctly predicted proteins in one of ten location classes. As additional measures tested (i.e., F1 score and Matthew correlation coefficient (MCC) (Gorodkin, 2004)) did not provide any novel insights, these were confined to the *Appendix: Additional Results*. Error estimates were calculated over ten random seeds on both test sets. For previous methods (DeepLoc and DeepLoc62 (Almagro Armenteros et al., 2017), LocTree2 (Goldberg et al., 2012), MultiLoc2 (Blum et al., 2009), SherLoc2 (Briesemeister et al., 2009), CELLO (Yu et al., 2006), iLoc-Euk (Chou et al., 2011), YLoc (Briesemeister et al., 2010) and WoLF PSORT (Horton et al., 2007)) published performance values were used (Almagro Armenteros et al., 2017) for *setDeepLoc*. For *setHARD*, the webserver for DeepLoc^1^ was used to generate predictions using either profile or BLOSUM inputs, whose results were later evaluated in Q10 and MCC. As a naive baseline, we implemented a method that predicted the same location class for all proteins, namely the one most often observed (in results: <u>Majority</u>). We provided code to reproduce all results^2^.

## 4. Results

### Embeddings outperformed MSAs

The simple EAT (embedding-based annotation transfer) already outperformed some advanced methods using MSAs (Fig. 2). The FNNs trained on *ProtT5* (Elnaggar et al., 2021) and ESM-1b (Rives et al., 2021) outperformed the SOTA *DeepLoc* (Almagro Armenteros et al., 2017) (Fig. 2). Methods based on *ProtT5* embeddings consistently reached higher performance values than other embedding-based methods (\**ProtT5* vs. rest in Figure 2). Results on Q10 were consistent with those obtained for MCC (*Appendix: Additional Results*).

**Figure 2.**
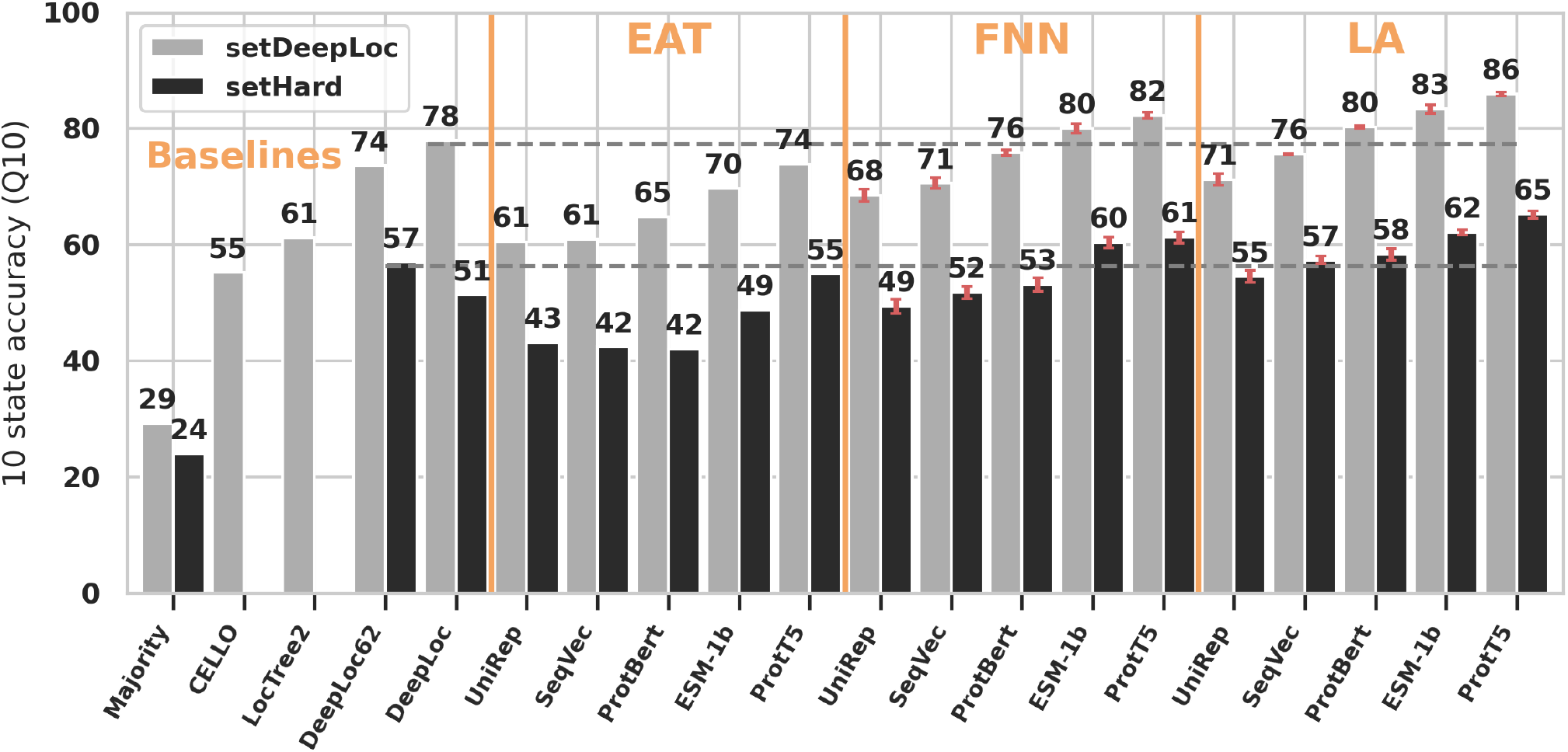
LA architectures performed best. <u>Performance</u>: Bars give the ten-class accuracy (Q10) assessed on *setDeepLoc* (light-gray bars) and *setHARD* (dark-gray bars). <u>Methods</u>: *Majority, CELLO*, LocTree2*, DeepLoc*, DeepLoc62;* MSA-based methods marked by star. <u>EAT</u> used the mean-pooled pLM embeddings to transfer annotation via distance, while <u>FNN(pLM)</u> used the mean-pooled embeddings as input to a feed-forward neural network. <u>LA(pLM)</u> marked predictions using light attention on top of the pLMs from: UniRep (Alley et al., 2019), SeqVec (Heinzinger et al., 2019), ProtBert (Elnaggar et al., 2021), ESM-1b (Rives et al., 2021), ProtT5 (Elnaggar et al., 2021). Horizontal gray dashed lines mark the previous SOTA (*DeepLoc* and *DeepLoc62*) on either set. Estimates for standard deviations are marked in red for the new methods. Overall, LA significantly outperformed the SOTA without using MSAs, and values differed substantially between the two data sets (light vs. dark gray).

### LA architecture best

The light attention (LA) architecture introduced here consistently outperformed other embedding-based approaches for all pLMs tested (LA* vs. EAT/FNN* in Fig. 2). Using *ProtBert* embeddings, LA outperformed the SOTA (Almagro Armenteros et al., 2017) by 1 and 2 percentage points on *setHARD* and *setDeepLoc* (*LA(ProtBert)* Fig. 2). For both test sets, LA improved the previous best on either set by around eight percentage points with *ProtT5* embeddings.

### Standard data set over-estimated performance

The substantial drop in performance measures (by about 22 percentage points) between the standard *setDeepLoc* and the new challenging *setHARD* (Fig. 2: light-gray vs. dark-gray, respectively) suggested substantial over-fitting. Mimicking the class distribution from *setDeepLoc* by sampling with replacement from *setHARD* led to higher values (Q10: DeepLoc62=63%; DeepLoc=54%; *LA(ProtBert)*=62%; *LA(ProtT5)*=69%). *DeepLoc* performed worse on *setHARD* with than without MSAs (only BLOSUM; Fig. 2: *DeepLoc* vs. *DeepLoc62*). Otherwise, the relative ranking and difference of models largely remained consistent between the two data sets *setDeepLoc* and *setHARD*.

### Low performance for minority classes

The confusion matrix of predictions for *setDeepLoc* using LA(*ProtT5*) highlighted how many proteins were incorrectly predicted to be in the second most prevalent class (*cytoplasm*), and that the confusion of the two most common classes mainly occurred between each other (Fig. 3: *nucleus* and *cytoplasm*). As for other methods, including the previous SOTA (Almagro Armenteros et al., 2017), performance was particularly low for the most under-represented three classes (*Golgi apparatus, lysosome/vacuole*, and *peroxisome*) that, together accounted for 6% of the data. To attempt boosting performance for minority classes, we applied a *balanced loss*, assigning a higher weight to the contributions of underrepresented classes. This approach did not raise accuracy for the minority classes but lowered the overall accuracy, thus it was discarded.

**Figure 3.**
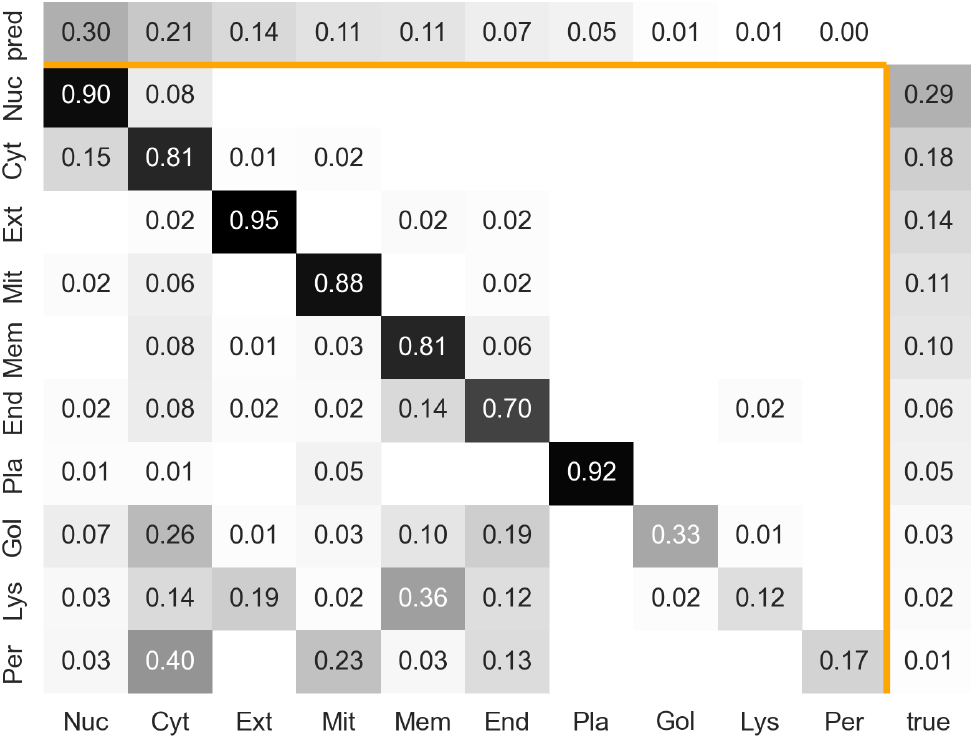
Mostly capturing majority classes. Confusion matrix of LA predictions on ProtT5 (Elnaggar et al., 2021) embeddings for *setDeepLoc* (Almagro Armenteros et al., 2017) (see *Appendix: Additional Results* for *setHARD*) Darker color means higher fraction; the diagonal indicates accuracy for the given class; vertical axis: true class; horizontal axis: predicted class. Labels are sorted according to prevalence in ground truth with the most common class first (left or top). Labels: Nuc=**Nuc**leus; Cyt=**Cyt**oplasm; Ext=**Ext**racellular; Mit=**Mit**ochondrion; Mem=cell **Mem**brane; End=**End**oplasmatic Reticulum; Pla=**Pla**stid; Gol=**Gol**gi apparatus; Lys=**Lys**osome/vacuole; Per=**Per**oxisome; **pred**=distribution for predicted (proteins predicted in class X / total number of proteins); **true**=distribution for ground truth (proteins in class X / total number of proteins)

### Light attention (LA) mechanism crucial

To probe the effectiveness of the LA aggregation mechanism on *ProtT5* we considered several alternatives for compiling the attention (LA *w/o Softmax* & *LA w/o MaxPool* & *Attention from v* & *DeepLoc LSTM* & *Conv* + *AdaPool*), and used the LA mechanism with non-embedding input (*LA(OneHot)* & *LA(MSA)*). Q10 dropped substantially without softmax- or max-aggregation. Furthermore, inputting *traditional* protein representations (*one-hot encoding*, i.e. representing the 20 amino acids by a 20-dimensional vector with 19 zeroes) or MSAs, the LA approach did not reach the heights of using pLM embeddings (Table 2: *LA(OneHot)* & *LA(MSA)*).

**Table 2.**
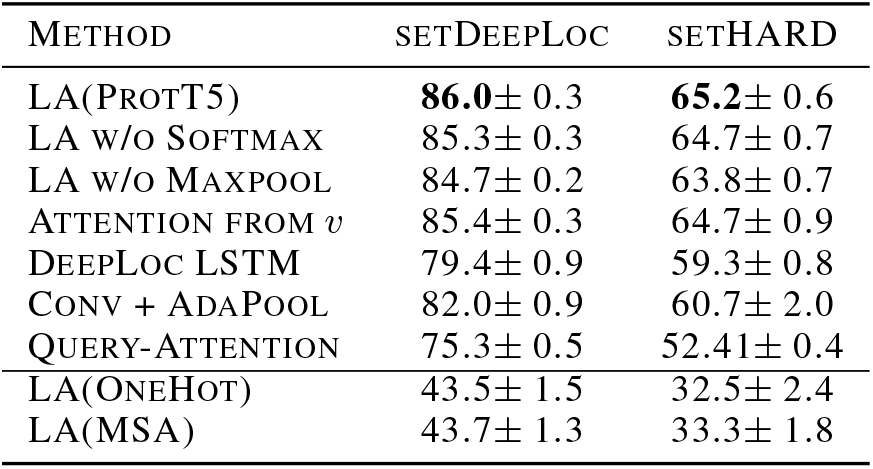
Comparison of <u>LA(ProtT5)</u> to different architectures and inputs. Methods described in Section 3.2. Standard deviations are estimated from 10 runs with different weight initializations.

### Model trainable on consumer hardware

Extracting ProtT5 pLM embeddings for all proteins used for evaluation took 21 minutes on a single Quadro RTX 8000 with 48GB vRAM. Once those input vectors had been generated, the final LA architecture, consisting of 19 million parameters, could be trained on an Nvidia GeForce GTX 1060 with 6GB vRAM in 18 hours or on a Quadro RTX 8000 with 48GB vRAM in 2.5 hours.

## 5. Discussion

### LA predicting location: beyond accuracy, four observations for machine learning in biology

The LA approach introduced here constituted possibly the largest margin to date of pLM embeddings improving over SOTA methods using MSAs. Although this improvement might become crucial to revive location prediction, ultimately this work might become even more important for other lessons learned:

i. The LA solution improved substantially over all previous approaches to aggregate per-residue embeddings into per-protein embeddings for predictions. Many protein function tasks require per-protein representations, e.g. predictions of Gene Ontology (<u>GO</u>), Enzyme Classifications (<u>E.C.</u>), binary protein-protein interactions (to bind or not), cell-specific and pathway-specific expression levels. Indeed, LA might help in several of these tasks, too.
ii. Although static, standard data sets (here the *DeepLoc* data) jumpstart advances and help in comparisons, they may become a trap for performance over-estimates through over-fitting. Indeed, the substantial difference in performance between *setDeepLoc* and *setHARD* highlighted this effect dramatically. Most importantly, our results underlined that claims of the type “*method NEW better than SOTA*” should not necessarily constitute wedges for advancing progress. For instance, *NEW* on *setStandard* reaching P(NEW)>P(SOTA) does not at all imply that *NEW* outperformed SOTA. Instead, it might point more to *NEW* over-fitting *setStandard*.
iii. The new data *setHARD* also pointed to problems with creating too well-curated data sets such as *setDeepLoc*: one aim in selecting a *good* data set is to use only the most reliable experimental results. However, those might be available for only some subset of proteins with particular features (e.g. short, well-folded). Experimental data is already extremely biased for the classes of location annotated (Marot-Lassauzaie et al., 2021). Cleaning up might even increase this bias and thereby limit the validity of prediction methods optimized on those data. Clearly, existing location data differ substantially from entire proteomes (Marot-Lassauzaie et al., 2021).
iv. *setHARD* also demonstrated that, unlike the protein structure prediction problem (Jumper et al., 2021), the location prediction problem remains unsolved: while Q10 values close to 90% for *setDeepLoc* might have suggested levels close to - or even above - the experimental error, *setHARD* revealed values of Q10 below 70%. In fact, while most proteins apparently mostly locate in one compartment, for others the multiplicity of locations is key to their role. This issue of *travellers vs. dwellers*, implies that Q10 cannot reach 100% as long as we count only one class as correctly predicted for each protein, and if we dropped this constraint, we would open another complication (Marot-Lassauzaie et al., 2021). In short, the new data set clearly generated more realistic performance estimates.

### Light attention (LA) beats pooling

The central challenge for the improvement introduced here was to convert the per-residue embeddings (NLP equivalent: word embeddings) from pLMs (*BB* (Bepler & Berger, 2019), *UniRep* (Alley et al., 2019), *SeqVec* (Heinzinger et al., 2019), *ProtBert* (Elnaggar et al., 2021), *ESM-1b* (Rives et al., 2021), and *ProtT5* (Elnaggar et al., 2021)) to meaningful per-protein embeddings (NLP equivalent: document). Qualitatively inspecting the influence of the light attention (LA) mechanism through a UMAP comparison (Fig. 4) highlighted the basis for the success of the LA. The embedding-based annotation transfer (EAT) surpassed some MSA-based methods without any optimization of the underlying pLMs (Fig. 2). In turn, inputting *frozen* pLM embeddings averaged over entire proteins into FNNs surpassed EAT and MSA-based methods (Fig. 2). The simple FNNs even improved over the SOTA, *DeepLoc*, for some pLMs (Fig. 2). However, LA consistently distilled more information from the embeddings. Most likely, the improvement can be attributed to LA coping better with the immense variation of protein length (varying from 30 to over 30 000 residues (Consortium, 2021)) by learning attention distributions over the sequence positions. LA models appeared to have captured relevant long-range dependencies while retaining the ability to focus on specific sequence regions such as beginning and end, which play a particularly important role in determining protein location for some proteins (Nair & Rost, 2005; Almagro Armenteros et al., 2017).

**Figure 4.**
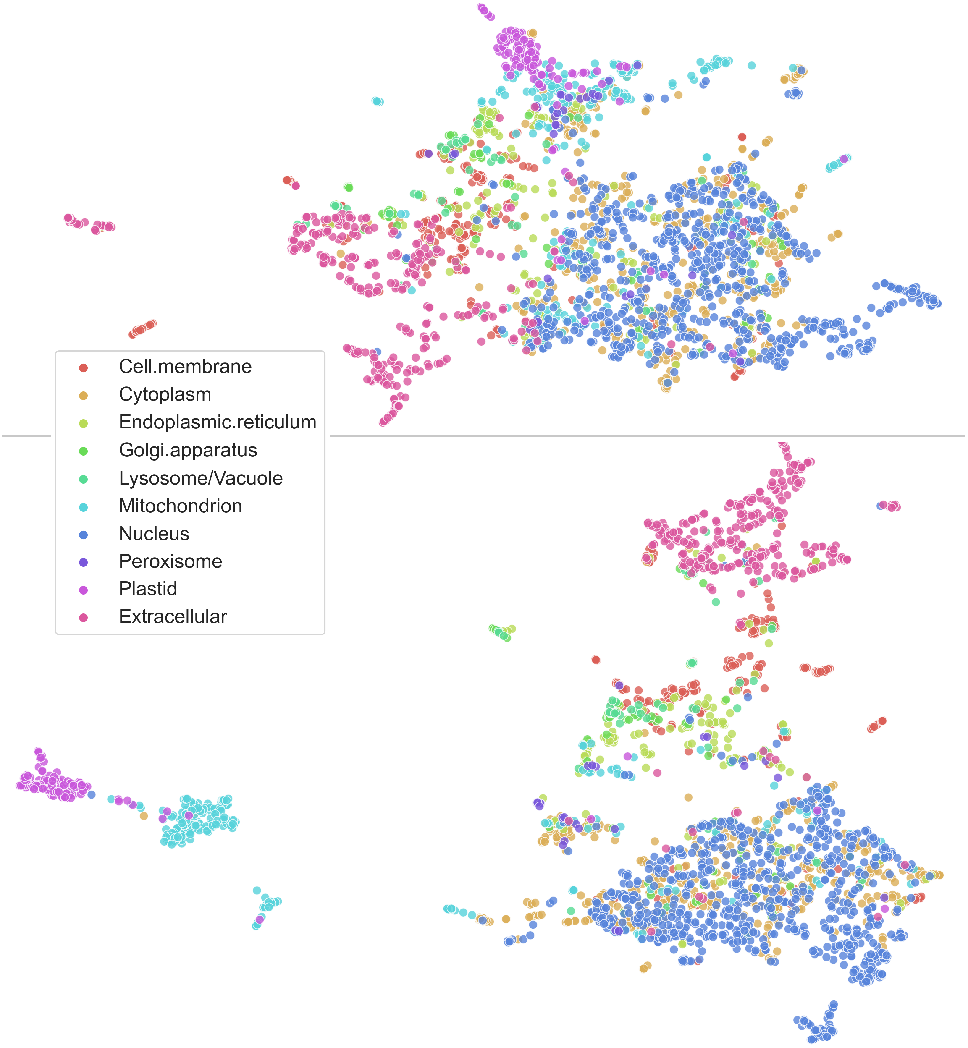
Qualitative analysis confirmed LA to be effective. UMAP (McInnes et al., 2018) projections of per-protein embeddings colored according to subcellular location (*setDeepLoc*). Both plots were created with the same default values of the python *umap-learn* library. Top: ProtT5 embeddings (LA input; *x*) mean-pooled over protein length (as for FNN/EAT input). Bottom: ProtT5 embeddings (LA input; *x*) weighted according to the attention distribution produced by LA (this is not *x*′ as we sum the input features *x* and not the values *v* after the convolution).

### Embeddings outperformed MSA: first for function

Effectively, LA trained on pLM embeddings from *ProtT5* (Elnaggar et al., 2021) was at the heart of the first method that clearly appeared to outperform the best existing method (*DeepLoc*, (Almagro Armenteros et al., 2017; Heinzinger et al., 2019)) in a statistically significant manner on a new representative data set not used for development (Fig. 2). To the best of our knowledge, it was also the first in outperforming the MSA-based SOTA in the prediction of subcellular location in particular, and of protein function in general. Although embeddings have been extracted from pLMs trained on large databases of unannotated (unlabelled) protein sequences that evolved, the vast majority of data learned originated from much more generic constraints informative of protein structure and function. Clearly, pre-trained pLMs never had the opportunity to learn protein family constraints encoded in MSAs.

### Better and faster than MSAs

When applying our solution to predicting location for new proteins (or *at inference*), the embeddings needed as input for the LA models come with three advantages over the historically most informative MSAs that were essential for methods such as *DeepLoc* (Almagro Armenteros et al., 2017) to become top. Most importantly, embeddings can be obtained in far less time than is needed to generate MSAs and require fewer compute resources. Even the lightning-fast MMseqs2 (Steinegger & Söding, 2017), which is not the standard in bioinformatics (other methods 10-100x slower), in our experience, required about 0.3 seconds per protein to generate MSAs for a large set of 10 000 proteins. One of the slowest but most informative pLMs (*ProtT5*) is three times faster, while the third most informative (*ProtBert*) is five times faster (Table 1). Moreover, these MMseqs2 stats derive from runs on a machine with >300GB of RAM and 2×40cores/80threads CPUs, while generating pLM embeddings required only a moderate machine (8 cores, 16GB RAM) equipped with a modern GPU with >7GB of vRAM. Additionally, the creation of MSAs relied on tools such as MMseqs2 that are sensitive to parameter changes, ultimately an extra complication for users. In contrast, generating embeddings required no parameter choice for users beyond the choice of the pLM (best here *ProtT5*). However, retrieving less specific evolutionary information (e.g. BLOSUM (Henikoff & Henikoff, 1992)) constituted a simple hash-table lookup. Computing such input could be instantaneous, beating even the fastest pLM *SeqVec*. Yet, these generic substitution matrices have rarely ever been competitive in predicting function (Ng & Henikoff, 2003; Bromberg et al., 2008). One downside to using embeddings is the one-off expensive pLM pre-training (Elnaggar et al., 2021; Heinzinger et al., 2019). In fact, this investment pays off if and only if the resulting pLMs are not retrained. If they are used unchanged - as shown here - the advantage of embeddings over MSA is increasing with every single new prediction requested by users (over 3,000/months just for *PredictProtein* (Bernhofer et al., 2021)). In other words, every day, embeddings save more over MSAs.

### Overfitting through standard data set?

For location prediction, the *DeepLoc* data (Almagro Armenteros et al., 2017) has become a standard. Static standards facilitate method comparisons. To solidify performance estimates, we created a new test set (*setHARD*), which was redundancy-reduced both with respect to itself and all proteins in the *DeepLoc* data (comprised of training plus testing data, the latter dubbed *setDeepLoc*). For *setHARD*, the 10-state accuracy (Q10) dropped, on average, 22 percentage points with respect to the static standard, *setDeepLoc* (Fig. 2). We argue that this large margin may be attributed to some combination of the following coupled effects.

i. Previous methods may have been substantially overfitted to the static data set, e.g., by misusing the test set to optimize hyperparameters. This could explain the increase in performance on *setHARD* when mimicking the class distributions in the training set and *setDeepLoc*.
ii. The static standard set allowed for some level of sequence-redundancy (information leakage) at various levels: certainly within the test set, which had not been redundancy reduced to itself (data not shown), maybe also between train and test set. Methods with many free parameters might more easily exploit such residual sequence similarity for prediction because proteins with similar sequences locate in similar compartments. In fact, this may explain the somewhat surprising observation that *DeepLoc* appeared to perform worse on *setHARD* using MSAs than the generic BLOSUM62 (Fig. 2: *DeepLoc62* vs. *DeepLoc*).Residual redundancy is much easier to capture by MSAs than by BLOSUM (Henikoff & Henikoff, 1992) (for computational biologists: the same way in which PSI-BLAST can outperform pairwise BLAST (Altschul et al., 1997)).
iii. The confusion matrix (Fig. 3) demonstrated how classes with more experimental data tended to be predicted more accurately. As *setDeepLoc* and *setHARD* differed in their class composition, even without overfitting and redundancy, prediction methods would perform differently on the two. In fact, this can be investigated by recomputing the performance on a similar class-distributed superset of *setHARD*, on which performance dropped only by 11, 24, 18, and 17 percentage points for *DeepLoc62, DeepLoc, LA(ProtBert)*, and *LA(ProtT5)*, respectively.

Possibly, several effects contributed to the performance from standard to new data set. Interestingly, different approaches behaved alike: both for alternative inputs from pLMs (*SeqVec, ProtBert, ProtT5*) and for alternative methods (EAT, FNN, LA), of which one (EAT) refrained from weight optimization.

### What accuracy to expect for the next 10 location predictions?

If the top accuracy for one data set was Q10 ~ 60% and Q10 ~ 80% for the other, what could users expect for their next ten queries: either six correct or eight, or between six and eight? The answer depends on the query: if those proteins were sequence similar to proteins with known location (case: redundant): the answer would be eight. Conversely, for new proteins (without homologs of known location), six in ten will be correctly predicted, on average. However, this assumes that the ten sampled proteins follow somehow similar class distributions to what has been collected until today. In fact, if we applied *LA(ProtT5)* to a hypothetical new proteome similar to existing ones, we can expect the distribution of proteins in different location classes to be relatively similar (Marot-Lassauzaie et al., 2021). Either way, this implies that for novel proteins, there seems to be significant room for pushing performance to further heights, possibly by combining *LA(ProtBert)/LA(ProtT5)* with MSAs.

## 6. Conclusion

We presented a light attention mechanism (LA) in an architecture operating on embeddings from several pLMs (*BB*, *UniRep, SeqVec, ProtBert, ESM-1b*, and *ProtT5*. LA efficiently aggregated information and coped with arbitrary sequence lengths, thereby mastering the enormous range of proteins spanning from 30-30 000 residues. By implicitly assigning a different importance score for each sequence position (each residue), the method succeeded in predicting protein subcellular location much better than methods based on simple pooling. More importantly, for three pLMs, LA succeeded in outperforming the SOTA without using MSA-based inputs, i.e., the single most important input feature for previous methods. This constituted an important breakthrough: although many methods had come close to the SOTA using embeddings instead of MSAs (Elnaggar et al., 2021), none had ever overtaken as the methods presented here. Our best method, *LA(ProtT5)*, was based on the largest pLM, namely on *ProtT5* (Fig. 2). Many methods were assessed on a standard data set (Almagro Armenteros et al., 2017). Using a new, more challenging data set (*setHARD*), the performance of all methods appeared to drop by around 22 percentage points. While class distributions and data set redundancy (or homology) may explain some of this drop, over-fitting might have contributed more. Overall, the drop underlined that many challenges remain to be addressed by future methods. For the time being, the best method *LA(ProtT5)* is freely available via a webserver (embed.protein.properties) and as part of a high-throughput pipeline (Dallago et al., 2021). Predictions for the human proteome are available via Zenodo https://zenodo.org/record/5047020.

## Acknowledgements

Thanks to Tim Karl (TUM) for help with hardware and software; to Inga Weise (TUM) for support with many other aspects of this work. Thanks to the Rostlab for constructive conversations and to the anonymous reviewers for constructive criticism. Thanks to all those who deposit their experimental data in public databases, and to those who maintain these databases. In particular, thanks to the Ioanis Xenarios (SIB, Univ. Lausanne), Matthias Uhlen (Univ. Upssala), and their teams at Swiss-Prot and HPA. This work was supported by the Deutsche Forschungsgemeinschaft (DFG) – project number RO1320/4-1, by the Bundesministerium für Bildung und Forschung (BMBF) – project number 031L0168, and by the BMBF through the program “Software Campus 2.0 (TU München)” – project number 01IS17049.

## Footnotes

1 http://www.cbs.dtu.dk/services/DeepLoc

2 https://github.com/HannesStark/protein-localization

