## Appendix for "Light Attention Predicts Protein Location from the Language of Life"

### A. Protein Preliminaries

**Protein Sequences.** Proteins are built by chaining an arbitrary number of one of 20 amino acids in a particular order. When amino acids come together to form protein sequences, they are dubbed residues. During the assembly in the cell, constrained by physiochemical forces, the one-dimensional chains of residues fold into unique 3D shapes based solely on their sequence that largely determine protein function. The ideal machine learning model would predict a protein's 3D shape and thus function from just protein sequence (the ordered chain of residues).

**Protein Subcellular Location.** Eukaryotic cells contain different organelles/compartments. Each organelle serves a purpose, e.g., ribosomes chain together new proteins while mitochondria synthesize ATP. Proteins are the machinery used to perform these functions, including transport in and out and communication between different organelles and a cell's environment. For some compartments, e.g., the nucleus, special stretches of amino acids, e.g., nuclear localization signals (NLS), help identifying a protein's location via simple string matching. However, for many others, the localization signal is diluted within the whole sequence, requiring sequence-level predictions. Furthermore, some organelles (and the cell itself) feature membranes with different biochemical properties than the inside or outside, requiring protein gateways.

**Homology-inference.** Two highly similar protein sequences will most likely fold in similar 3D structures and are more likely to perform similar functions. Homology-based inference (Nair & Rost, 2002; Mahlich et al., 2018), which transfers annotations of experimentally validated proteins to query protein sequences, is based on this assumption (Sander & Schneider, 1991). Practically this means searching a database of annotated protein sequences for sequences that meet both an identity threshold and a length-of-match threshold to some query protein sequence. Sequence homology delivers good results, but its stringent requirements render it applicable to only a fraction of proteins (Rost, 1999).

**Machine learning Function Prediction.** When moving into territory where sequence similarity is less conserved for shorter stretches of matching sequences (Mahlich et al., 2018; Rost, 2002), one can try predicting function using evolutionary information and machine learning (Goldberg et al., 2012; Almagro Armenteros et al., 2017). Evolutionary information from protein profiles, encoding a protein's evolutionary path, is obtained by aligning sequences from a protein database to a query protein sequence and computing conservation metrics at the residue level. Using profiles leads to impressively more accurate predictions for sequences with no close homologs and has been the standard for most protein prediction tasks (Urban et al., 2020), including subcellular localization (Goldberg et al., 2012; Almagro Armenteros et al., 2017; Savojardo et al., 2018). While profiles provide a strong and useful inductive bias, their information content heavily depends on a balance of the number of similar proteins (depth), the overall length of the matches (sequence coverage), the diversity of the matches (column coverage), and their generation is parameter sensitive.

### B. Hyperparameters

The following describes the search space used to find hyperparameters of our final LA and FNN models. We performed random search over these parameters. The evaluated learning rates were in the range of $[5 \times 10^{-6}$ - $5 \times 10^{-3}]$. For the light attention architecture, we tried filter sizes $[3, 5, 7, 9, 11, 13, 15, 21]$ and hidden sizes $d_{out} \in [32, 128, 256, 512, 1024, 1500, 2048]$, as well as concatenating outputs of convolutions with different filter sizes. For the FNN, we searched over the hidden layer sizes $[16, 32, 64, 512, 1024]$, where $32$ was the optimum. We maximized batch size to fit a Quadro RTX 8000 with 48GB vRAM, resulting in the batch size of $150$. Note that the memory requirement is dependent on the size of the longest sequence in a batch. In the DeepLoc dataset, the longest sequence had 13 100 residues.

### C. Datasets

Since ESM-1b can only process sequences shorter than 1024 residues, we removed the longer ones. This resulted in 8662 sequences for the training data, 2457 for *setDeepLoc*, and 431 for *setHard*. Table 3 shows the distribution of subcellular localization classes in the standard *setDeepLoc* and our new *setHARD* with all sequences included.

*Table 3.* Number of proteins and percentage of dataset for each class for the DeepLoc dataset and our *setHARD*. ER abbreviates Endoplasmatic Reticulum

| LOCATION | DEEPLOC | | SETHARD | |
|---|---|---|---|---|
| | # | % | # | % |
| NUCLEUS | 4043 | 28.9 | 99 | 20.2 |
| CYTOPLASM | 2542 | 19.3 | 117 | 23.8 |
| EXTRACELLULAR | 1973 | 14.0 | 92 | 18.8 |
| MITOCHONDRION | 1510 | 11.8 | 10 | 2.0 |
| CELL MEMBRANE | 1340 | 9.5 | 98 | 20.0 |
| ER | 862 | 6.2 | 34 | 6.9 |
| PLASTID | 757 | 5.4 | 11 | 2.6 |
| GOLGI APPARATUS | 356 | 2.6 | 13 | 2.6 |
| LYSOSOME/VACUOLE | 321 | 2.3 | 13 | 2.2 |
| PEROXISOME | 154 | 1.1 | 3 | 0.6 |

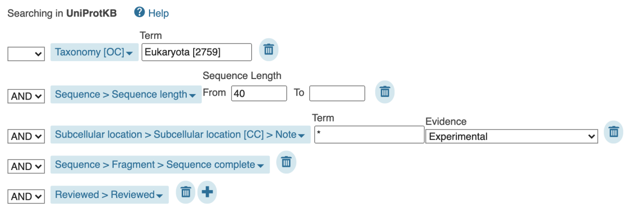


*Figure 5.* Screenshot of the filtering options applied to the advanced UniProt search (uniprot.org/uniprot).

##### C.1. New test set creation

In the following, we lay out the steps taken to produce the new test set (*setHARD*). The starting point is a filtered UniProt search with options as selected in Figure 5. Python code used is available at data.bioembeddings.com/public/data/new_test_set_procedure_code_data.zip.

- Download data as FASTA & XML:

```
wget "https://www.uniprot.org/
uniprot/?query=taxonomy:%
22Eukaryota%20[2759]%22%
20length:[40%20TO%20*]%
20locations:(note:*%20evidence:%
22Inferred%20from%20experiment%
20[ECO:0000269]%22)%20fragment:no%
20AND%20reviewed:yes&format=
xml&force=true&sort=score&compress=
yes"
```

```
wget "https://www.uniprot.org/
uniprot/?query=taxonomy:%
22Eukaryota%20[2759]%22%
20length:[40%20TO%20*]%
20locations:(note:*%20evidence:%
22Inferred%20from%20experiment%
20[ECO:000026%22)%20fragment:no%
20AND%20reviewed:yes&format=
fasta&force=true&sort=
score&compress=yes"
```

- Download deeploc data:

```
wget http://www.cbs.dtu.dk/services/
DeepLoc-1.0/deeploc_data.fasta
```

- Align sequences in swissprot to deeploc that have more than 20% PIDE:

```
mmseqs easy-search swissprot.fasta
deeploc_data.fasta -s 7.5
--min-seq-id 0.2 --format-output
query,target,fident,alnlen,mismatch,
gapopen,qstart,qend,tstart,tend,
evalue,bits,pident,nident,qlen,tlen,
qcov,tcov alignment.m8 tmp
```

- Extract localizations from SwissProt XML:

```
python extract_localizaiotns
_from_swissprot.py
```

- Map deeploc compartments on swissprot localizations & remove duplicates ([P123, Nucleus] appearing twice), remove multilocated ([P123, Nucelus] and [P123, Cytoplasm] –> remove P123) empty or not experimental annotations:

```
python map_and_filter
_swissprot_annotations.py
```

- Create FASTA like deeploc from sequences not in alignment:

```
python extract_unaligned_sequences.py
```

- Redundancy reduce new set to 20%:

```
mmseqs easy-cluster
--min-seq-id 0.2 new_test_set
_not_redundancy_reduced.fasta
new_hard_test_set_PIDE20.fasta tmp
```

#### D. Additional Results

We provide results for both *setDeepLoc* (Table 7) and *setHARD* (Table 6) in tabular form, including the Matthew's Correlation Coefficients (MCC) and class unweighted F1 score.

Furthermore, in Figure 6 we find that the UMAP projections of $x'$ are more similar to those of the attention coefficients pooled along the length dimension and show clear clusters. Meanwhile, the projections of $v^{max}$ in Figure 7 are less informative even though the ablations showed that $v_m ax$ is important for the performance of our architecture.

Notable is that for both projections there are some clear outliers with the localization Plastid that are mapped far away from all other projections.

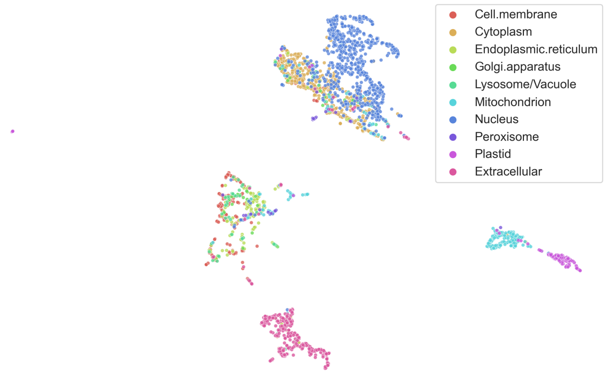


*Figure 6.* UMAP (McInnes et al., 2018) projections of $x'$ embeddings colored according to subcellular location (*setDeepLoc*).

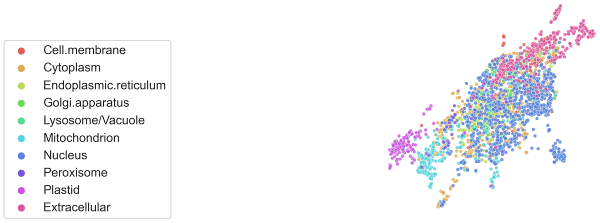


*Figure 7.* UMAP (McInnes et al., 2018) projections of $v^{max}$ embeddings colored according to subcellular location (*setDeepLoc*).

*Table 4.* MCC of additional baselines and ablations compared to the LA architecture on *ProtT5* embeddings (above the line) of *setDeepLoc* and *setHARD* averaged over 10 seeds. The best method is **bold** and the second best is underlined.

| METHOD | SETDEEPLOC | SETHARD |
|---|---|---|
| LA PROTT5 | **.831**± .004 | **.577**± .007 |
| LA - SOFTMAX | .828± .004 | .570± .008 |
| LA - MAXPOOL | .816± .002 | .559± .008 |
| ATTENTION FROM V | .824± .003 | .571± .012 |
| DEEPLOC LSTM | .752± .010 | .505± .009 |
| CONV + ADAPOOL | .785± .010 | .526± .022 |
| MEANPOOL + FFN | .785± .006 | .529± .010 |
| LA ON ONEHOT | .326± .012 | .216± .014 |
| LA ON PROFILES | .302± .016 | .195± .022 |

*Table 5.* Class unweighted F1 score of additional baselines and ablations compared to the LA architecture on *ProtT5* embeddings (above the line) of *setDeepLoc* and *setHARD* averaged over 10 seeds. The best method is **bold** and the second best is underlined.

| METHOD | SETDEEPLOC | SETHARD |
|---|---|---|
| LA PROTT5 | **.854**± .004 | **.642**± .004 |
| LA - SOFTMAX | .850± .004 | .633± .008 |
| LA - MAXPOOL | .842± .002 | .632± .006 |
| ATTENTION FROM V | .845± .004 | .634± .011 |
| DEEPLOC LSTM | .788± .009 | .590± .007 |
| CONV + ADAPOOL | .818± .010 | .608± .020 |
| MEANPOOL + FFN | .814± .005 | .604± .008 |
| LA ON ONEHOT | .367± .025 | .262± .033 |
| LA ON PROFILES | .384± .018 | .279± .019 |

*Table 6.* Accuracy and Matthew's correlation coefficient (MCC) on *setHard*. Baseline= predict majority class; Evo= Previous methods based on evolutionary inputs; AT= assign class based on nearest neighbour in embedding space; FNN= predict using a Multi-Layer Perceptron on top of embeddings; LA= predict using LA on top of embeddings; Embedding inputs from: BB (Bepler & Berger, 2019), UniRep (Alley et al., 2019), SeqVec (Heinzinger et al., 2019), ProtBert (Elnaggar et al., 2021), ESM-1b (Rives et al., 2021), ProtT5 (Elnaggar et al., 2021).

| | Method | Accuracy | MCC |
|---|---|---|---|
| | Baseline | 24 | 0 |
| Evo | DeepLoc62 | 56.94 | 0.476 |
| | DeepLoc | 51.36 | 0.410 |
| AT | BB | 25.98 | 0.133 |
| | UniRep | 43.15 | 0.329 |
| | SeqVec | 42.43 | 0.315 |
| | ProtBert | 42.04 | 0.306 |
| | ESM-1b | 48.72 | 0.386 |
| | ProtT5 | 55.01 | 0.454 |
| FNN | BB | 35.60± 2.34 | 0.247± 0.025 |
| | UniRep | 49.41± 1.21 | 0.391± 0.013 |
| | SeqVec | 51.71± 1.04 | 0.398± 0.013 |
| | ProtBert | 53.16± 1.19 | 0.429± 0.014 |
| | ESM-1b | 60.40± 0.94 | 0.518± 0.010 |
| | ProtT5 | 61.27± 0.97 | 0.529± 0.010 |
| LA | BB | 40.80± 2.44 | 0.293± 0.027 |
| | UniRep | 54.56± 1.07 | 0.451± 0.011 |
| | SeqVec | 57.37± 0.64 | 0.468± 0.013 |
| | ProtBert | 58.36± 1.02 | 0.490± 0.012 |
| | ESM-1b | 62.12± 0.5 | 0.537± 0.004 |
| | ProtT5 | **65.21**± 0.61 | **0.577**± 0.007 |

*Table 7.* Accuracy and Matthew's correlation coefficient (MCC) on *setDeepLoc*. Baseline= predict majority class; Evo= Previous methods based on evolutionary inputs; AT= assign class based on nearest neighbour in embedding space; FNN= predict using a Multi-Layer Perceptron on top of embeddings; LA= predict using LA on top of embeddings; Embedding inputs from: BB (Bepler & Berger, 2019), UniRep (Alley et al., 2019), SeqVec (Heinzinger et al., 2019), ProtBert (Elnaggar et al., 2021), ESM-1b (Rives et al., 2021), ProtT5 (Elnaggar et al., 2021).

| | Method | Accuracy | MCC |
|---|---|---|---|
| | Baseline | 29 | 0 |
| Evo | LocTree2 | 61.20 | 0.525 |
| | MultiLoc2 | 55.92 | 0.487 |
| | SherLoc2 | 58.15 | 0.511 |
| | YLoc | 61.22 | 0.533 |
| | CELLO | 55.21 | 0.454 |
| | iLoc-Euk | 68.20 | 0.641 |
| | WoLF PSORT | 56.71 | 0.479 |
| | DeepLoc62 | 73.60 | 0.683 |
| | DeepLoc | 77.97 | 0.735 |
| AT | BB | 40.94 | 0.295 |
| | UniRep | 60.54 | 0.519 |
| | SeqVec | 60.97 | 0.508 |
| | ProtBert | 64.85 | 0.567 |
| | ESM-1b | 69.67 | 48.72 |
| | ProtT5 | 73.92 | 0.687 |
| FNN | BB | 48.43± 0.99 | 0.367± 0.011 |
| | UniRep | 68.49± 1.02 | 0.622± 0.011 |
| | SeqVec | 70.57± 0.93 | 0.636± 0.011 |
| | ProtBert | 75.88± 0.45 | 0.702± 0.006 |
| | ESM-1b | 80.02± 0.84 | 0.760± 0.009 |
| | ProtT5 | 82.28± 0.51 | 0.786± 0.006 |
| LA | BB | 55.75± 0.89 | 0.462± 0.010 |
| | UniRep | 71.24± 0.96 | 0.654± 0.011 |
| | SeqVec | 75.63± 0.11 | 0.705± 0.002 |
| | ProtBert | 80.29± 0.21 | 0.762± 0.002 |
| | ESM-1b | 83.39± 0.76 | 0.8013± 0.009 |
| | ProtT5 | **86.01**± 0.34 | **0.832**± 0.004 |

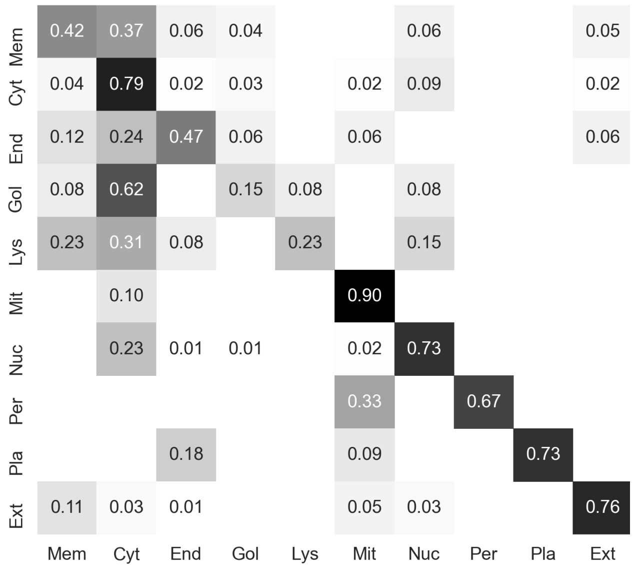


*Figure 8.* Confusion matrix of LA predictions on ProtT5 (Elnaggar et al., 2021) embeddings for *setHARD* annotated with the fraction of the true class. Vertical axis: true class, horizontal axis: predicted class. Labels: Mem=cell **Mem**brane; Cyt=**Cyt**oplasm; End=**End**oplasmatic Reticulum; Gol=**Gol**gi apparatus; Lys=**Lys**osome/vacuole; Mit=**Mit**ochondrion; Nuc=**Nuc**leus; Per=**Per**oxisome; Pla=**Pla**stid; Ext=**Ext**racellular
